# Multi –omics and metabolic modelling pipelines: challenges and tools for systems microbiology

**DOI:** 10.1101/013532

**Authors:** Marco Fondi, Pietro Liò

## Abstract

Integrated -omics approaches are quickly spreading across microbiology research labs, leading to i) the possibility of detecting previously hidden features of microbial cells like multi-scale spatial organisation and ii) tracing molecular components across multiple cellular functional states. This promises to reduce the knowledge gap between genotype and phenotype and poses new challenges for computational microbiologists. We underline how the capability to unravel the complexity of microbial life will strongly depend on the integration of the huge and diverse amount of information that can be derived today from - omics experiments. In this work, we present opportunities and challenges of multi –omics data integration in current systems biology pipelines. We here discuss which layers of biological information are important for biotechnological and clinical purposes, with a special focus on bacterial metabolism and modelling procedures. A general review of the most recent computational tools for performing large-scale datasets integration is also presented, together with a possible framework to guide the design of systems biology experiments by microbiologists.

## Introduction

The ease at which genomes are currently sequenced has assigned to genomics one of the first steps in microbial systems biology. Regardless of the technique used, assembly and annotation typically follow genome sequencing and return an almost complete picture of the genetic reservoir of a given microorganism. On the other hand, genome sequence only represents a snapshot of the real phenotypic capabilities of an organism, providing very few indications on other crucial aspects of the underlying life cycle such as response to environmental and genetic perturbations, fluctuations in time, gene essentiality and so on. To gain a systemic and exhaustive description of living entities, static information deriving from genome sequence is not enough and other levels of knowledge must be taken into consideration. Nowadays, technologies do exist for measuring, in a large-scale fashion, other crucial aspects of cellular life, including the level of RNA within the cell (transcriptomics), the nature of metabolites present within the cell (metabolomics), the interaction among different proteins (protein-protein interaction) and many others (detailed below). Also, metabolic biodiversity of microbial communities can be today evaluated through metagenomics and metatranscriptomics approaches. However, no single –omics analysis can fully unravel the complexities of fundamental microbiology (Zhang et al., 2010). Multi- and integrated -omics approaches have thus started spreading among several research areas, from bio-based fuel production (Zhu et al., 2013) to biopharmaceuticals processes (Schaub et al., 2012), from medical research (Wiench et al., 2013) to host-pathogen interactions (Ansong et al., 2013b). The integration of such diverse data types may be considered one of the key challenges of present-day bioinformatics, due to different data formats, high data dimensionality and need for data normalization.

One of the most important drawbacks associated with the booming of genomics resides in the possibility to (almost) automatically derive the potential metabolic landscape of a strain, given its genome. Bacteria continuously provide industry with novel products/processes based on the use of their metabolism and numerous efforts are being undertaken to deliver new usable substances of microbial origin to the marketplace (Beloqui et al., 2008), including pharmaceuticals, biofuels and bioactive compounds in general (George et al., 1983; Garcia-Ochoa et al., 2000; Lee et al., 2005; Zou et al., 2012). In this context, computational modelling and *in silico* simulations are often adopted by metabolic engineers to quantitatively simulate chemical reactions fluxes within the whole microbial metabolism. To exploit computational approaches, genome annotation-derived metabolic networks are transformed into models by defining the boundaries of the system, a biomass assembly reaction, and exchange fluxes with the environment (Durot et al., 2009). Also needed are i) structured (mathematical) representation of that network, ii) possibly quantitative parameters enabling simulations or predictions on the joint operation of all network reactions in a given environment and, in particular, iii) predictions on the values of metabolite fluxes and/or concentrations (Papin et al., 2003). A constraint-based modelling framework can then be used to automatically compute the resulting balance of all the chemical reactions predicted to be active in the cell and, in turn, to bridge the gap between knowledge of the metabolic network structure and observed metabolic processes (Varma and Palsson, 1994).

Innovative high-throughput technologies (see Figure 1) represent a valuable resource also in the context of metabolic modelling, since data integration can be performed to gain a clearer and more comprehensive picture of the metabolic traits of a given organism. Diverse data types can be mapped onto metabolic models in order to elucidate more thoroughly the metabolism of a cell and its response to environmental factors; this is usually done by including functional characterization and accurate quantification of all levels of gene products, mRNA, proteins and metabolites, as well as their interaction (Zhang et al., 2010).

**Figure 1.**
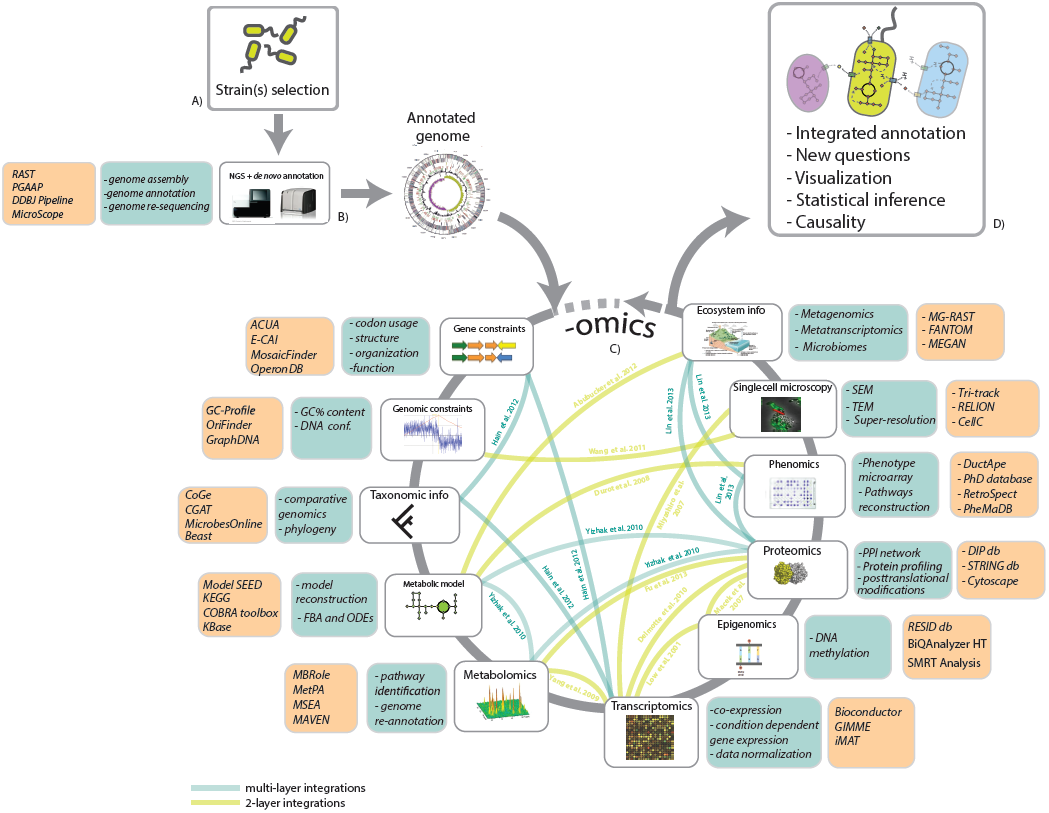
Data integration in microbial –omics pipelines. A: Strain isolation and preliminary experimental characterization usually leads to the identification/selection of bacteria that may be of potential interest in biotechnology. B: Nowadays, to gain a systems level perspective on biotechnologically relevant strains NGS and preliminary genome annotation is usually performed. After this step, information on the presence/absence of metabolic pathways and overall metabolic capabilities of a given microbe is gained. C: Nevertheless, to obtain a systems-level knowledge, a body of additional information can be mapped onto a genome annotation (the –omics wheel). This includes: gene and genomic constraints (derived from a deep inspection of genome properties), taxonomic and metabolic information, –omics data (transcriptomics, proteomics, metabolomics, epigenomics, phenomics), other phenotypic information (e.g. high-resolution microscopy), ecosystem information, (microbiome composition, community functional characterization, meta-transcriptomics). Furthermore, these different layers of information can be combined and integrated to merge together datasets resulting from the application of different technologies. Links among –omics represent present-day study cases in which integration among two or more information layers has been performed (see corresponding references). D: After –omics integration has been performed, a more comprehensive perspective on the microbe(s) under study is gained, providing clues on the possible interactions with the surrounding environment (including metabolic cross-talk with other microbial species), statistically grounded inferences and novel questions to be addressed (possibly re-iterating the pipeline). In this figure, orange boxes include possible software (see Supplementary Table 2 for the full list) while green-blue boxes specific tasks of general –omics strategy.

Here we review and discuss possible experimental and computational pipelines for multiple data integration in microbial research, allowing the simultaneous analysis of different data-types and their mapping onto a *de novo* genome annotation. We discuss which layers of biological information have been shown to be important for biotechnological/clinical purposes and whether these layers are independent or have to be considered as a single complex system. Computational insights will be reviewed, including data mining, pre-processing, assimilation and iterative integration in order to exploit all available information.

Furthermore, given the link existing between microbial phenotypes and underlying metabolism, we will discuss a general framework of the major steps and checkpoints encountered when reconstructing the metabolic network of a given organism and in its consequent exploitation for computational simulation and/or phenotype prediction.

We underline the importance of integrating different sources of information to gain a more comprehensive view of genome annotation and metabolic features in general.

## Information layers overview

This section provides a schematization of the sources of information (layers) that are currently the most exploited in systems biology. These layers represent the basis of multi-omics integration discussed in the next section.

### Microbial genomics pipelines

Since fast genome sequencing and preliminary data post-processing have been achieved, well-grounded experimental design and strains selection have (re)gained a key position when drafting genomics-oriented research plans (Fig. 1A). Complete genome sequence is increasingly more often the starting point for integrative pipelines (see below and Fig. 1). Sample preparation and sequencing can be performed in a few days while data postprocessing (including quality check, *de novo* assembly, gene prediction) still represents the most demanding bottleneck of the genomics pipeline. For this reason, most of the leading genomics centres (including BGI, DOE-JGI, Craig Venter Institute, Sanger) couple genome sequencing to bioinformatics analysis and usually make their software open access to the research community. In this way, preliminary assembly, annotation and analysis of genomes are carried out, although many other popular tools exist for *de novo* genome annotation (see for example (Angiuoli et al., 2008; Aziz et al., 2008; Seemann, 2014)).

Besides basic genomic knowledge (e.g. gene presence/absence patterns), many other additional information layers are today available to be merged and integrated when trying to fully elucidate complex biological patterns of living entities (Fig. 1). References to study cases and computational tools related to these informational layers are reported in Supporting Information 1 and 2, respectively. These include:

- **Gene constraints** represent the additional information present in DNA sequences and not fully exploited by functional annotation pipelines. These may include the detection of gene fusions, the identification of operons and the computation of gene Codon Adaptation Index (CAI). Gene structure data (e.g. the presence of gene fusions) can guide the identification of potential protein–protein interactions (Enright et al., 1999). Further, the study of operonic organization exploits co-directional intergenic distances and can provide homology-free functional annotation through the transfer of functions among co-operonic genes (i.e. genes belonging to the same operon) through the so-called the ‘guilt by association’ principle. Lastly, CAI measures the variability of the codon usage in a gene in respect to the variability of a reference set of genes and usually provides a reliable index of potential protein expression levels within a cell.
- **Genomic constraints** include compositionally heterogeneous G+C domains in DNA sequences that can provide much insight into biological features such as horizontal gene transfer (HGT) events, pathogenicity, and antibiotic resistance. Also, 3D structure of chromosomes is known to regulate many DNA-based processes including repair, replication, and transcription.
- **Metabolic information** (the repertoire of metabolic reactions possessed by a microbe) can be automatically computed starting from genome annotations. Derived metabolic reconstructions (models) can be exploited for *in silico* metabolic modelling and simulation, for example using so called constraints-based methods (e.g. flux balance analysis, FBA) that are currently widely adopted (mainly because they do not require detailed information on the chemical equations of the studied system).
- **Phylogenetic information** refers to the potential represented by the huge taxonomic range of sequenced genomes available in public database. More specifically, it involves the use/integration of pieces of information extrapolated from closely related organisms to fill knowledge gaps of the microbe(s) under investigation.
- **Epigenomics** can be today investigated at single-base and strand resolution and is gaining a central role since sometimes it can explain phenotypic differences arising among cells with identical genetic information. DNA methylation state of particular regions, for example, is linked to chromosomes partitioning, gene expression and virulence and DNA repair. Unlike eukaryotes, bacteria use DNA adenine methylations as epigenetic signals (rather than DNA cytosine methylation). Similar to eukaryotes, however, spatial organization of phenotypes was recently shown to be controlled and/or regulated by an **epigenetic memory** effect in bacteria (Kurz et al., 2013), thus establishing a link between metabolism, ecology and tuning cellular phenotype. Importantly, links between epigenetic mechanisms and bacterial pathogenesis/virulence have been established (Casadesus and Low, 2006; Shell et al., 2013).
- **Metabolomics** provides the metabolic profiling of a given organism in the form of an instantaneous snapshot of the physiology of that cell. Mapping the outputs metabolomic technologies onto genome annotations may improve the accuracy of existing gene models metabolic pathways identification and genomes re-annotations
- **Transcriptomics** (generated and measured through high-density tiling microarrays or transcriptome shotgun sequencing using next generation technology) can yield an improved description of metabolic processes active in well-defined environmental conditions, also providing useful functional insights since many functionally related genes (e.g. belonging to the same metabolic pathway) have been shown to be co-expressed.
- **Proteomics** refers to the study of proteins structure, function and abundance at the whole-cell level. Accordingly, it allows the quantitation of the level of expression of cellular proteins; also it can shed light onto post-translational modifications [that can be identified, for example, by surface plasmon resonance (SPR) combined with Mass Sprectometry (MS), named SPR-MS (Buijs and Franklin, 2005), by glycoproteomics pipelines (Hitchen and Dell, 2006) or by a plethora of possible phosphoproteomics approaches (Lin et al., 2010)], protein-protein interaction (PPI, detected, for example, by means of co-immunoprecipitation. See (Rao et al., 2014) for a review) and protein localization (for example studied by immunofluorescence and fluorescent-protein tagging (Stadler et al., 2013)). PPI data, for example, have proven valuable for inferring protein function from functions of interaction partners. Additionally, protein phosphorylation can alter protein functions by changing structure/function relationship as well as enzymes catalysis/interaction and allowing a quick adjustment to changing environment.
- **Phenomics** (high-throughput phenotyping) allows exploring the phenotypic space of a given organism and deriving phenotype capabilities (e.g. the capability of metabolizing certain carbon sources in respect to others) in an automated and large-scale fashion. This is typically performed through Phenotype MicroArrays (PMs) (Bochner et al., 2001) which uses cellular respiration (i.e. NADH reduction) and consequent production of a purple colour as a reporter system for overall metabolic activity.
- **Microscopy** can provide high quality imaging of microbial life and detailed knowledge on a number of cellular components, including i) protein expression, localization and interaction (through fluorescence and confocal laser scanning microscopy), ii) alterations of bacterial cell envelope and interaction between host and bacterial cells (Transmission and Scanning Electron Microscope, TEM and SEM, respectively) and iii) detection of single molecules (through super-resolution microscopy, SRM (Schermelleh et al., 2010; Vogelsang et al., 2010)).

## Integration of information layers

To date, many examples of multi-layer studies exist in scientific literature (Fig. 1C), ranging from simpler integrations (two different –omics datasets) to more comprehensive and computationally demanding ones (multiple –omics data).

Two layers integration include approaches that combine **transcriptomics and metabolomics** (Kromer et al., 2004; Durre, 2007; Yoshida et al., 2008; Depuydt et al., 2009; Yang et al., 2009; Sana et al., 2010), **proteomics and metabolomics** (Ma et al., 2011; Fu et al., 2013) and **proteomics and transcriptomics** (Huang et al., 2013). These latter kind of studies, in particular, have shown how gene expression regulation can be largely decoupled from protein dynamics and how translation efficiency can a have higher regulatory impact on protein abundance than protein turnover (Jayapal et al., 2008; Maier et al., 2011). These recent studies suggest a relevant role of post-transcriptional, translational and degradation regulation in determining cellular protein concentrations and that they may contribute at least as much as transcription itself (Vogel and Marcotte, 2012).

An interesting two layer integration was recently proposed (Burton et al., 2014) to reconstruct the individual genomes of microbial species present within a mixed sample and combining metagenomics with chromatin-level contact probability maps [generated with the Hi-C method (Lieberman-Aiden et al., 2009)]. By applying this approach to synthetic metagenomes data, authors succeeded in clustering genome content of fungal, bacterial, and archaeal species with great accuracy (99% agreement with published reference genomes)

Other, less exploited integrated approaches, comprise i) the integration of **metatranscriptomics and metagenomics** for determining the functional role of each microorganism in relation to the composition of the microbial community it is inserted into (Shi et al., 2011), ii) the integration of **comparative genomics and transcriptomics** (Hain et al., 2012) and iii) the **combination of photo-activated localization microscopy and proteomics** (Endesfelder et al., 2013)

Three different information layers (encompassing **metabolomics**, **transcriptomics** and **genomics**) were combined with metabolic modeling of *Mycoplasma pneumoniae* (Maier et al., 2013). Results obtained led the authors to infer a certain rigidity of metabolic pathway architecture in *M. pneumoniae*, suggesting that these are regulated as functional units rather than on the level of individual enzymatic reactions probably allowing a simplification of metabolic fluxes adjustment (Maier et al., 2013). Deatherage et al. (2013) applied a combined approach (which included **proteomics**, **metabolomics**, **glycomics**, **and metagenomics**) and identified complex metabolic interplay among the intestinal microbiome including the capability of S. *enterica* to suppress the growth of Bacteroidetes and Firmicutes representatives while promoting growth of *Salmonella* and *Enterococcus* ones (Deatherage Kaiser et al., 2013).

**Transcriptomics, proteomics and translatomics** (the evaluation of mRNAs in polysome fractions) were combined by Berghoff et al. (2013) in order to evaluate the dynamic and regulatory features of bacterial oxidative stress responses of the purple bacterium *Rhodobacter sphaeroides*, leading to the creation of a multi-layered expression map on the system level (expressome) (Berghoff et al., 2013). Authors found that, in this case, translational control appears to exceed simple regulation at the transcriptional level and that gene positioning might be involved in the tuning of the expression patterns within inducible operons. Finally, (Perco et al., 2010) recently proposed the integration of several -omics profiles from similar datasets at the level of PPI, thus including a further level of information in these kind of studies.

Super-meta approaches (Fig. 2) are the result of these multidimensional integrations, capable of combining multiple heterogeneous information layers. Also, from these examples [and (Poblete-Castro et al., 2012; Ansong et al., 2013a; Chang et al., 2013; Karaosmanoglu et al., 2013; Zhu et al., 2013)] it emerges that multi-omics, same condition, experiments promise to be crucial in the future of systems microbiology. Indeed, since complex phenotypes may arise as a result of the action of different biological processes (e.g. transcription, translation, post-translational modifications and so on) an integrated multi –omics approach may reveal which of them is (or are) contributing the most to the observed cellular behaviour.

### Computational aspects of information layers integration

In large scale, multi-omics studies, the computational resources allocation needed for data processing and integration quickly outpaces the resource allocation for data generation (Palsson and Zengler, 2010) and one of the greater efforts resides in analysing the outcome of a set of experiments in a unified and interactive way.

In a broader sense, integration of different information layers poses (at least) three main computational challenges, that is i) tracking the different molecular components (i.e. gene, molecules, enzymes) across different datasets and experiments (Task 1), ii) identifying (or developing) reliable data normalization procedures and multi-level data analysis (Task 2) and iii) producing an effective visualization of the results (Task 3).

**Task 1.** The first task is achieved by linking gene names to codes that are shared across multiple databases during genome annotation. This is usually done on the basis of orthology relationships across gene datasets and sequence DBs, although other pieces of information can provide hints for refining functional classification (gene structure/organization, see below). COG (Tatusov et al., 2003), KAAS (Moriya et al., 2007) and GO (Dimmer et al., 2012) are widely adopted tools for functional annotation that allow the retrieval of gene-centred information during multi-layer study. Similarly, databases such as ChEBI (Hastings et al., 2013), PubChem (Li et al., 2010) provide unique identifiers for linking the same compound across multiple datasets or experiments.

**Task 2.** Concerning the second step (normalization procedures and multi-layer data analysis), linear regression (Park et al., 2003), central tendency (global normalization) (Yang et al., 2002) and singular value decomposition (Karpievitch et al., 2009) are widely adopted techniques to reduce systematic errors in *n*-high-throughput studies. However, since different –omics layers may possess characteristic rates of noise, variation or discrepancies caused by complex molecular mechanisms, accurate normalization procedures are required when performing statistical analyses over a multi-layer dataset (Arakawa and Tomita, 2013). The Expression Index (EI), for example, accounts for the global change within a specific type of cellular component measured by the corresponding analytical method. Focused statistical analyses on EI values across multi-layer datasets enable highlighting differences between samples in various perturbation experiments (Ishii et al., 2007; Arakawa and Tomita, 2013).

Tools have been developed to facilitate handling and correlating large and diverse datasets under the same working environment and allowing measuring a number of different metrics (like pathway overrepresentation, inter-associations between pathways and diseases, enrichment analyses) (Zhang and Drabier, 2012; Sun et al., 2013). A list of currently available computational tools for downstream analysis and biological interpretation of omics data is presented in Table 1. Although with differences among each other, these tools allow tracing the relationships existing between different biological functional levels (e.g. gene expression, metabolism) and their corresponding molecular components (e.g. genes, metabolites) by integrating results from two (or more) –omics experiments. MONA is a model-based Bayesian tool for inferring significant associations across multiple information layers such as mRNA and protein expression profiling as well as DNA methylation and microRNA regulation (Sass et al., 2013a). Besides providing a pipeline for data analysis of different high-throughput approaches, MADMAX database also tackles the issue of data storage in multi-omics experiments (Lin et al., 2011). CONFERO can store the lists of genes derived from **a priori** biological knowledge and integrate them with results from -omics data. Statistics on these multi-level datasets can then be performed including, for example, functional enrichment analyses. By exploiting a method called Kriging (typical of geostatistics and machine learning), Omickriking computes a genomic similarity among the samples using a linear combination of -omics similarity matrices and predicts the emergence of cellular complex traits through weighted average of the phenotype of individuals in the training set. VANTED provides a framework for mapping and integrating experimental data over biochemical networks that are drawn by the user or downloaded from the KEGG database. NetGestalt is a tool suited for presenting multi-scale experimental data and facilitating their integration, visualization and analysis. Each information layer is represented on a separate track but this structure can be easily turned into a network that can be visualized using the implemented Cytoscape web plug-in (Shannon et al., 2003). Also, six different methods are available for analysing data stored in the tracks, such as, for example, the identification of enriched network modules or enrichment in GO terms or metabolic pathways.

**Table 1.** List of software/methods for–omics integration

| Method/Software | Website | Ref. |
| --- | --- | --- |
| **-omics datasets integration** |  |  |
| OmicKriging (R package) | http://cran.r-project.org/web/packages/OmicKriging/index.html | - |
| Integromics | https://www.integromics.com/ | (Le Cao et al., 2009) |
| MONA | http://sourceforge.net/projects/monaos/ | (Sass et al., 2013b) |
| NetGestalt | http://www.netgestalt.org/ | (Shi et al., 2013) |
| VANTED | http://vanted.ipk-gatersleben.de/ | (Klukas and Schreiber, 2010) |
| mixOmics (R package) | http://www.qfab.org/mixomics/ | - |
| Confero | http://sourceforge.net/projects/confero/ | (Hermida et al., 2013) |
| DAnTE | http://omics.pnl.gov/software/DanteR.php | (Polpitiya et al., 2008) |
| MADMAX | http://madmax2.bioinformatics.nl:443/dev/f?p=104:1 | (Lin et al., 2011) |
| MCIA | http://www.bioconductor.org/packages/release/bioc/html/omicade4.html | (Meng et al., 2014) |
| InCroMAP | http://www.ra.cs.uni-tuebingen.de/software/InCroMAP/ | (Wrzodek et al., 2013) |
| **Causality** |  |  |
| qtlnet | http://cran.r-project.org/web/packages/qtlnet/ | (Neto et al., 2010) |
| pcalg | http://pcalg.r-forge.r-project.org/ | (Markus Kalisch, 2012) |
| **-omics data visualization** |  |  |
| OmicsAnalyzer | http://apps.cytoscape.org/apps/omicsanalyzer | (Xia et al., 2010) |
| GALANT | http://www.lbbc.ibb.unesp.br/galant/ | (Camilo et al., 2013) |
| BisoGenet | http://apps.cytoscape.org/apps/bisogenet | (Martin et al., 2010) |
| CytoHiC | http://www.cl.cam.ac.uk/~ys388/CytoHiC/ | (Shavit and Lio, 2013) |
| MODAM | http://www.softpedia.com/get/Programming/Components-Libraries/MODAM.shtml | (Nagy et al., 2013) |
| NuChart | ftp://fileserv.itb.cnr.it/nuchart | (Merelli et al., 2013) |
| Pathway Tools | http://bioinformatics.ai.sri.com/ptools/ | (Paley and Karp, 2006) |
| ProMeTra | https://prometra.cebitec.uni-bielefeld.de/cgi-bin/prometra.cgi?login=prometra | (Neuweger et al., 2009) |
| Paintomics | http://bioinfo.cipf.es/node/826 | (Garcia-Alcalde et al., 2011) |
| 3Omics | http://3omics.cmdm.tw/ | (Kuo et al., 2013) |
| Metscape | http://metscape.ncibi.org/ | (Karnovsky et al., 2012) |
| MAYDAY | http://www-ps.informatik.uni-tuebingen.de/mayday/ | (Battke et al., 2010) |

InCroMAP (Wrzodek et al., 2013) currently supports the simultaneous analysis of mRNA, miRNA (microRNA), DNA methylation and protein (modification) data. By means of a hypergeometric test InCroMAP is able to assess the significance of the overrepresentation of predefined gene sets in multiple platforms experiments. Results obtained with this tool and representing the structured view of the metabolic changes present in a certain experimental condition can be interactively browsed and visualized by means of the classical KEGG representation scheme.

The R-package (http://www.R-project.org) provides a set of tools for multi-omics integration. DAnTE, for example, performs statistical and quantitative analyses of different high-throughput datasets that can be imported in the format of a simple *csv* file (Polpitiya et al., 2008). Moreover it provides a graphical front-end for basic data analysis tasks in –omics disciplines, although being particularly suited for proteomics datasets. Similarly, IntegrOmics can perform correlation analysis among (-omics) variables measured for the same sample and provided in the form of two-block data matrices.

A multivariate approach to the integration of multi-omics datasets has been recently proposed (named MCIA, Multiple Co-Inertia Analysis) (Meng et al., 2014). This method allows the identification of co-relationships between multiple high dimensional datasets through an exploratory data analysis method and was shown to fit both heterogeneous (proteomics and transcriptomics) and homogeneous (microarray and RNAseq based transcriptomics) datasets. An important characteristic of this tool resides in the fact that, since it does not rely on genomic feature annotation, it is not limited by the well-known issue of the incompleteness of present day annotations. Finally, mixOmics provides statistical integrative techniques [regularized Canonical Correlation Analysis (CCC) and sparse Partial Least Square, (PLS)] to analyse highly dimensional and heterogeneous data sets and to unravel relationships them.

Another consequence that stems for the collection of multiple, standardised, highly-dimensional 'omics' datasets from living organisms resides in the possibility to investigate causal relationships between genomics differences, phenotypes and -omics datasets in general. Granger causality has been applied to multi-omics study cases (Walther et al., 2010; Doerfler et al., 2013) and tools have been developed for identifying cause-effect relationships between experimental variables in large scale datasets (Table 1).

An alternative strategy may consist in performing multi-layer data integration through a 2 step network approach and in considering, from a statistical perspective, the -omics data point as a huge space of causal models. Indeed, multiple -omics data sets can be represented in the form of a multiple-weighted network where vertex and vertex weights represent –omics data whereas data regarding functional or physical interactions between components are represented as edges and edge weights. Multi Objective (MO) optimisation could be then used to find networks that are optimal according to several criteria, corresponding to desired phenotypes. These criteria can be implemented as functions of the biological data. For example, these functions (objective functions) can be defined to calculate overrepresentation analysis statistics among datasets. Optimisation quality indicators, such as the hyper volume, are used to establish comparisons among different runs of MO optimisation. Important directions of methodological development will be the multi parameter evidence synthesis and methods for causality inference based on extensions of sparse instrumental variable approaches (Forbes and Griffiths, 2002; Carlo Berzuini, 2012).

**Task 3.**, Finally, when analysing different layers of information, it is useful to produce an integrated view of the obtained results and represent quantitative data of the complete cascade from genotype to phenotype for individual organisms. Available methods (see (Gehlenborg et al., 2010) for an exhaustive review) can be roughly divided in i) tools for automated, network-oriented representation and analysis of large biological datasets and ii) tools focused on assembly and curation of pathways. Network visualization is often adopted in this context since it is effective in representing diverse (cellular) components and the interactions existing among them and providing intuitive interpretation of multilayer data. Cytoscape is probably the most popular tool for visualizing biological networks. Through a large set of plugins it permits specific analyses and different combinations of – omics integration also allowing researchers to map multiple omics datasets over the same cellular components network (Table 1).

Mapping heterogeneous –omics data using customized metabolic pathway maps is also a common practice and a large plethora of alternatives are currently available (Table 1). The Pathway Tools offers an overview diagram of the biochemical networks and pathways of an organism including the possibility to combine and represent gene expression and metabolomics measurements over it (Paley and Karp, 2006). The web-based ProMeTra visualizes and integrates datasets from several –omics datasets on user defined metabolic pathway maps (Neuweger et al., 2009). Similarly, Paintomics is able to derive lists of significant gene or metabolite changes from transcriptomics and metabolomics data and paint this information on KEGG-derived metabolic maps (Garcia-Alcalde et al., 2011). 3Omics software is a web based tool able to perform inter-omics correlation analyses and to visualize relationships in data with respect to time, experimental conditions and data types (Kuo et al., 2013). Experimental data for metabolites, genes and pathways can be combined and integrated in the context of relevant metabolic networks with the software Metscape (Karnovsky et al., 2012). Finally, MAYDAY is an application for analysis and visualization of general -omics data and providing functional insights with the possibility to combine gene expression, metabolomes and biosynthetic pathways and visualize results as differentially coloured pathway diagrams (Battke et al., 2010).

## Metabolic modelling

Given the link existing between phenotypic features of a given microbe and its underlying metabolism, in this part of the review we provide a general overview of the major steps and checkpoints encountered when reconstructing the metabolic network of a given organism and in its consequent exploitation for computational simulation and/or phenotype prediction (Fig. 3). We will here focus on stoichiometric rather than kinetic modelling. A kinetic reaction network consists of biochemical reactions that can be traditionally described by ordinary or partial differential equations (ODEs and PDEs, respectively) (Tomar and De, 2013). The application of kinetic models, however, is limited to small (well characterized) biochemical systems since many intracellular experimental measurements are needed in this modelling framework. Conversely, stoichiometric (constraints based) modelling can be applied to larger (genome scale) biochemical systems since it requires only the information on metabolic reactions stoichiometry and mass balances around the metabolites under pseudo-steady state assumption (Oberhardt et al., 2009).

Interestingly, global-scale properties of metabolic networks have been inferred using network modelling techniques. These include, for example, the emergence of bow-tie structures (Friedlander, 2014), latent versatility and carbon efficiency (Bardoscia, 2014), robustness and plasticity of metabolic pathways (Berger et al., 2014), identification of synthetic lethal reaction sets in metabolic networks, essential plasticity and redundancy of metabolism (Guell et al., 2014). The application of metabolic modelling approaches seems promising also in the context of the emerging issue of antibiotic resistance, as it has been shown that specific metabolic traits are crucial for the persistence of multi-drug resistant microbial sub-populations [the so-called persisters, (Prax and Bertram, 2014)]. Finally, as new genomes from the same taxonomic unit (e.g. bacterial species) are being released, comparative metabolic modelling approaches are being exploited for studying the correlation between unique strain-specific metabolic capabilities and their corresponding pathotypes and/or environmental niches (Monk et al., 2013).

Currently available computational tools for *in silico* metabolic modelling have been recently reviewed in (Copeland et al., 2012; Moura et al., 2013) and will not be covered here.

### A metabolic modelling pipeline

The input of a common metabolic modelling flowchart is usually a (complete or draft) genome sequence (generally referred to as “NGS” in Fig. 3A). Publically available resources exist for automatically generating draft metabolic reconstructions (see Supporting Information 2). Draft metabolic models usually require extensive gap-filling for the identification of metabolic functions that have not been identified during preliminary reconstruction step; this can usually be performed using comparative genomics approaches using highly curated models of (more or less distantly) related microorganisms (see next section). Once gaps have been correctly identified and filled, a first comparison between phenotypes prediction capabilities of the model and experimental data (i.e. experimentally validated growth rates) can be performed (Fig. 3B). Initial reconstruction usually fails in correctly identifying all the metabolic capabilities, and extensive model refinement is required for reconciling model predictions with phenotypic data (e.g. Biolog data, Fig. 3C). When a fitting is found between experimental data and *in silico* predictions, well-grounded computational simulation can start (Fig. 3D), according to the biological rationale of the overall metabolic modelling project [e.g. optimisation of biomass production and overproduction of a given compound using, for example, Pareto optimality and multiobjective functions (Angione et al., 2013; Sengupta et al., 2013)]. Outcomes of computational simulations typically require experimental validation; in case this is achieved, the reconstruction process can generally be considered terminated and the model can be used for further *in silico* analyses (e.g. knockouts simulation, robustness analysis, changes in growth medium composition, etc.) or to suggest further wet-lab experiments. Importantly, a working metabolic model can be used for the refinement of other metabolic models (possibly from closely related microorganisms), thus connecting the end of the pipeline with the beginning stages of reconstruction of other organisms (i.e. C and A boxes of Fig. 3).

**Figure 2.**
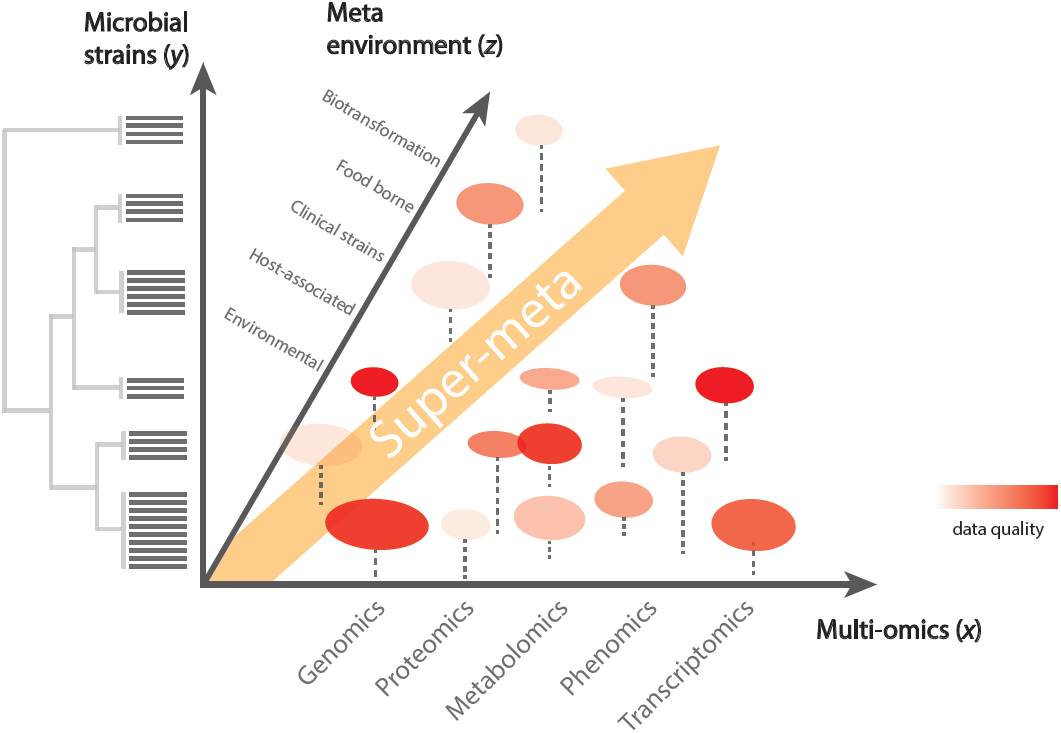
Super-meta and multi-parameter evidence synthesis approaches for –omics integration. Available technologies and public datasets allow approaching systems biology issues through a super-meta approach. Accordingly, different combinations of microbial ensembles (*y* axis), sampling environments (*z* axis) and/or –omics technologies (*x* axis) can be integrated and analysed, exploiting, for example multi evidence synthesis approaches. In this figure, the different circles represent different datasets, with hypothetical data quality represented in red-scale and the amount of data by circles size.

**Figure 3.**
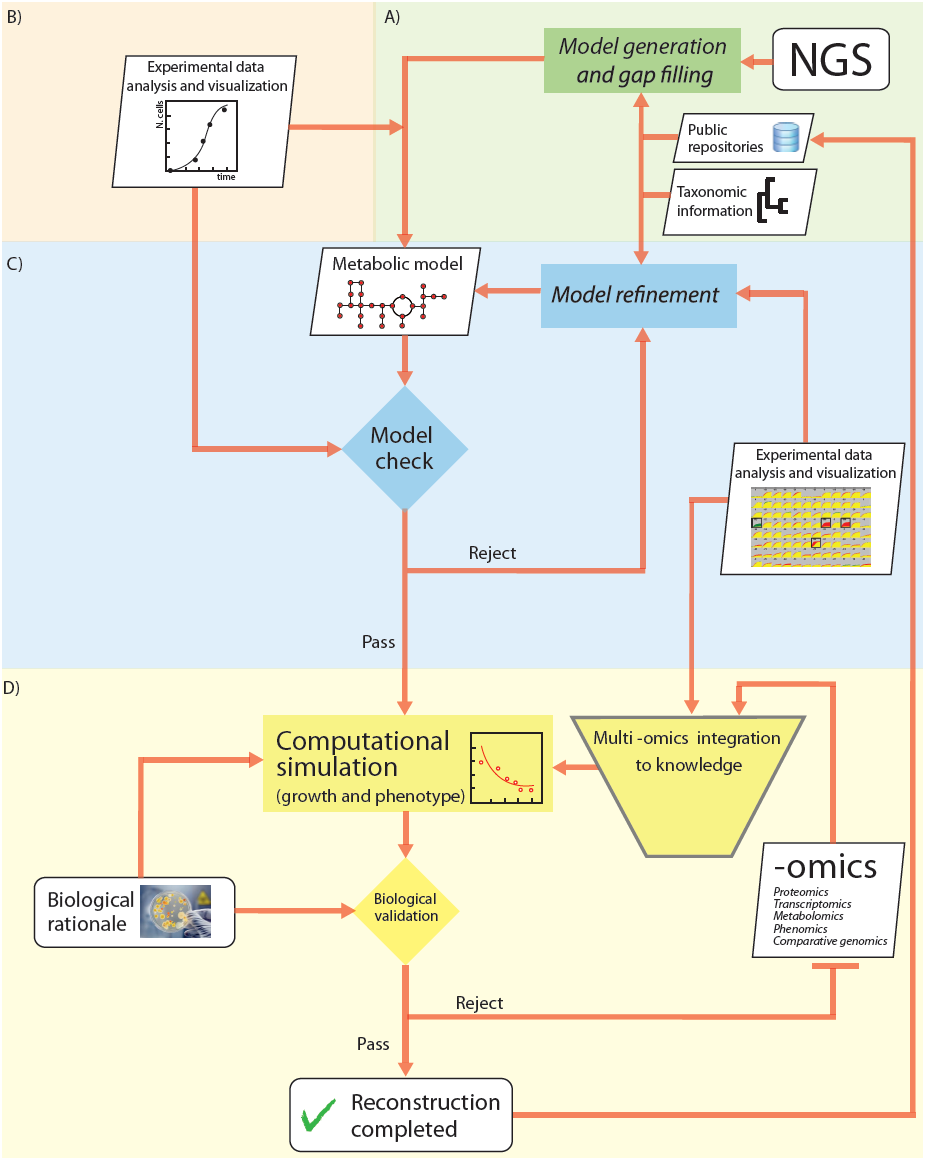
Overall scheme of a metabolic reconstruction/modelling pipeline and possible – omics integration. A: The input is typically a genome resulting from NGS. Tools for automatically generating draft metabolic reconstructions from public repositories are reported in Supplementary Table 1). B: Experimental data (i.e. growth rates, phenotype microarrays) can be used at this moment to help model construction/refinement. C: Draft metabolic models usually undergo extensive gap-filling. Once gaps have been correctly identified and filled, a first comparison between phenotypes prediction capabilities of the model and experimental data. D: When a fitting is found between experimental data and *in silico* predictions, more grounded computational simulation start. In this phase, –omics data integration (e.g. transcriptomics, proteomics) can be used for refining and reconciling modelling predictions. Finally, a working and experimentally validated metabolic model can be used for the refinement of other metabolic models.

Finally, in case no agreement is found or a greater accuracy is required, in this final stage additional (experimental) data (typically -omics data, -omics box” in Fig. 3D) should be produced in order to integrate, fix and expand the reconstructed model. This integration is also useful when a given (metabolic) phenotype cannot be explained relying exclusively the metabolic information layer.

### Which –omics information could improve metabolic modelling?

Despite fine-scale cellular features - such as molecular crowding (Beg et al., 2007) and macromolecular synthesis (Thiele et al., 2012) - can be accounted for by metabolic modelling, –omics-derived data is often used to assist and improve metabolic model predictions and to provide a system-level understanding of the cellular behaviour (in Fig. 3D).

Comparative genomics/taxonomy between multiple organisms can be exploited during metabolic reconstructions especially when gap-filling a newly reconstructed model or when trying to include novel reactions into it. Indeed, as each metabolic reconstruction is a reflection of the genetic content of the respective organism, the identification of the overlaps between the genomic content of the organisms for which a metabolic reconstruction is available can help understanding which reaction should be added/removed to/from a network. The refinement of the metabolic models from two *Pseudomonas* representatives is an example such an approach (Oberhardt et al., 2011). Full comparative analysis of their genome-scale metabolic reconstructions has led, in this case, to the identification of differences in their virulence mechanisms and in their metabolic features that, in turn, may be of help for future metabolic engineering application.

Transcriptomics is one of the most exploited layers for improving or correcting metabolic predictions. By exploiting gene expression data, genome scale metabolic networks can be turned into *condition specific* models in which only those reactions corresponding to expressed genes will be present and active [see for example (Fondi et al., 2014)]. In practice, this corresponds to respectively “turn on” or “turn off” those reactions whose corresponding genes are up- or down-regulated *in vivo*, when running the *in silico* simulation.

When mapping gene expression data onto metabolic models, data derived from multicondition, single platform –omics studies are of value, in that results from every single experiment can be easily compared with those from the other replicates (Faith et al., 2007; Colijn et al., 2009). In some cases, such approach has been shown to provide a realistic picture of the actual metabolic state of a microbial cell and to lead to a deeper understanding of its basic functioning, including the consequences of environmental perturbations such as gene knock-outs and/or growth medium manipulation.

Similarly, under the assumption that protein abundance changes correlate to changes in metabolic fluxes, quantitative proteomics data can be used for deriving condition-specific metabolic models.

Relative proteomics (and transcriptomics) data can be useful for studying system-level changes of the metabolism following an external perturbation (e.g. changes in growth temperature, medium, pH, etc.). In a recent work, for example, we have mapped protein abundance data onto the newly reconstructed metabolic model of the Antarctic bacterium *Pseudoalteromonas haloplanktis* TAC125 (Fondi et al., 2014). This has led to the identification of biologically consistent metabolic adjustments caused by cold shock-dependent changes in protein expression. Similarly, metabolic network modelling and proteome data were combined in (Tong et al., 2013) to explore the metabolic consequences of a downshift in culture temperature in the anaerobic thermophilic bacterium *Thermoanaerobacter tengcongensis* and to decipher the effect of proteome change on the bacterial growth under perturbation.

Absolute quantitation of protein abundance, instead, can be used for *de novo* drafting of metabolic models(Thomas et al., 2014; Vanee et al., 2014). Indeed, the use of high-throughput proteomics data as a starting point proved to be the most accurate in resembling in vivo growth conditions (Vanee et al., 2014).

Several methods are today available for integrating gene expression with (constraints-based) modelling (see Table 1 and (Blazier and Papin, 2012)). GIMME and iMAT maximise the flux across those reactions whose genes display high mRNA levels and minimise those of reactions associated with low mRNA levels according to a user-supplied expression threshold. MADE does not require an *a priori* threshold on expression levels, relying on multiple expression datasets to account for differential expression and constraint fluxes across model reactions. Unlike the above-mentioned approaches, E-FLUX provides more physiologically consistent solutions since it does not convert expression levels to binary states (active or inactive reactions). Besides constraints based modelling and expression data, PROM allows integrating pre-compiled transcription regulatory networks. GX-FBA combines hierarchical regulation imposed by gene-expression with the constraint of metabolic reaction connectivity and using information on mRNA levels to guide hierarchical regulation of metabolism subject to the interconnectivity of the metabolic network. Finally, RELATCH takes advantage of the concept of relative optimality (based on relative flux changes with respect to a reference flux distribution) and uses flux and gene expression data to predict metabolic responses in a genetically or environmentally perturbed state. Remarkably, the application of all these methods may be hampered by the observation that, in some cases, mRNA transcript levels do not correlate with protein levels (Gygi et al., 1999; Nie et al., 2006). In those cases, other information layers (e.g. post-translational modifications, post-transcriptional regulation) may explain this discrepancy and, in turn, their integration during metabolic modelling procedures might provide more realistic insights and/or predictive capability. A systematic evaluation of the available methods for integrating transcriptomics data into constraint-based models of metabolism has been recently performed (Machado and Herrgard, 2014), revealing that none of the methods outperforms the others for all the tested conditions. Also, in most cases, the predictions obtained by simple FBA using growth maximization and parsimony [i.e. assuming that the cell attempts to achieve its objective while allocating the minimum amount of resources (Machado and Herrgard, 2014)] criteria still represents a preferable choice for a realistic picture of the metabolic landscape of the organism under study.

Stoichiometric network modelling can be integrated with *in vivo* measurements of metabolic fluxes to determine the absolute flux through large networks of the carbon metabolism, using FBA, ^13^C fluxomics or ^13^C-constrained FBA approaches (Winter and Kromer, 2013). Directly comparing the result of an *in silico* simulation with isotopologue data, for example, can help in addressing which of the predicted metabolic fluxes is real and how far its value is from the *in vivo* measurements. Alternatively, ^13^C-based metabolic flux analysis (^13^C-MFA) can be formulated as an inverse problem to compute a set of fluxes that leads to the best match of the experimentally measured fluxes.

For example, a ^13^C-MFA network model was generated for *E. coli* revealing the reliability of this integrated approach to predict and measure the operation and regulation of metabolic networks (Chen et al., 2011). Similarly, by applying ^13^C metabolic flux analysis and *in silico* FBA, insights into xylose metabolism in *S. cerevisiae* were obtained, including futile pathways and the link between high cell maintenance energy and xylose utilization (Feng and Zhao, 2013).

A hypothesis-driven algorithm for the integration of transcriptomics and metabolomics data with metabolic network topology was originally developed and its application to the metabolic network of *Saccharomyces cerevisiae* showed the feasibility of inferring whether biochemical reactions within a cell are hierarchically or metabolically regulated (Cakir et al., 2006). A novel method (IOMA) has been developed to quantitatively integrate proteomic and metabolomic data with genome-scale metabolic models using a mechanistic model for determining reaction rate (Yizhak et al., 2010). The integration of such sources of information has led to achieve a greater accuracy in predicting the metabolic state of *E. coli* under different gene knockouts in respect to other methods (i.e. FBA and MOMA, <u>M</u>inimization <u>O</u>f <u>M</u>etabolic <u>A</u>djustment).

*In silico* modelling growth predictions on defined media can be easily compared with Biolog substrate utilization data (phenomics). This is typically achieved by (qualitatively) comparing the *estimated* flux value across biomass assembly reaction of the model with the activity directly measured during phenotype microarray experiment. In other words, growth predictions on a given sole carbon source (either growth or no growth) are compared against the outcome of a large scale phenotype profiling experiment and in case no agreement is found some reconciliation steps are usually necessary. These include, for example, the inclusion of a transport reaction for a specific tested compound (for which disagreement of *in silico* and *in vivo* data is observed) or the inclusion of previously missing metabolic reactions (gap-filling).

This comparison usually speeds up the identification of missing transport reactions and/or metabolic gaps in the model and, in some cases, can give essential functional insights concerning unknown genes. A number of currently available metabolic reconstructions have been validated adopting such approach, including models of *Burkholderia cenocepacia* J2315 (Fang et al., 2011), *Acinetobacter baylyi* ADP1 (Durot et al., 2008), *Bacillus subtilis* (Oh et al., 2007). Most of the metabolic reconstructions that are currently available display accuracy values ranging from 75% up to more than 95% when their growth predictions are compared against phenotype microarrays growth data.

A list of available computational tools for integrating –omics datasets and metabolic models is reported in Table 2.

**Table 2.**
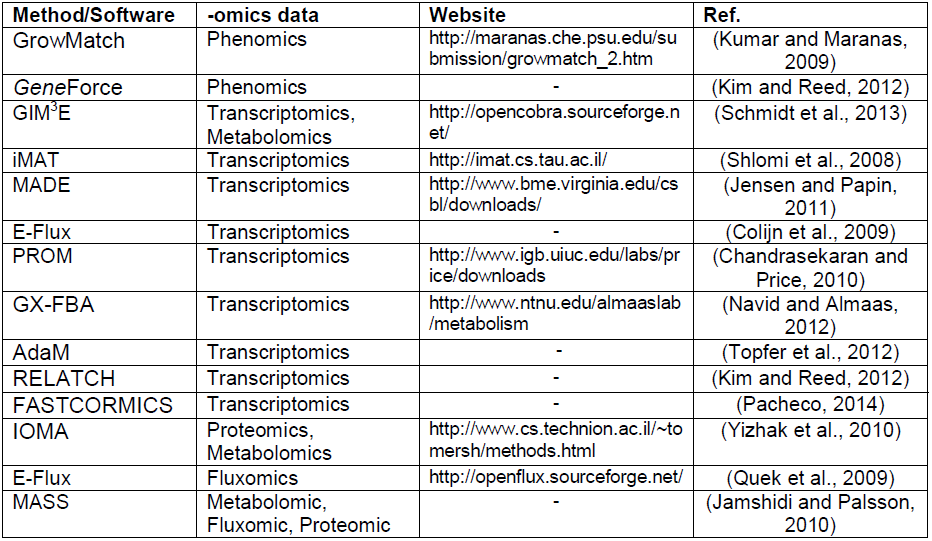
List of software/methods for integrating –omics data and metabolic modeling

### What is missing in metabolic reconstruction?

Currently, about 4000 complete genome sequences are available in public databases (www.genomesonline.org); conversely, only around 100 reconstructions of microbial metabolic systems can be retrieved (see http://systemsbiology.ucsd.edu/InSilicoOrganisms/OtherOrganisms for an updated list). This gap is the most evident consequence of the difficulties in reconstructing “working” metabolic models starting from genome annotations and speeding up this process is a key challenge for future systems microbiology (Fig. 2).

Also, the sketchiness of present-day metabolic models is an important point that needs consideration. Indeed, even in the case of model organisms, reconstructions rarely account for any more than roughly 25% of their genes, leading to an underestimation of their real metabolic capabilities of the strain under study. However, in some cases, such an apparently small fraction of genes embedded in metabolic reconstructions may indeed represent the majority of the metabolic genes that are actually expressed during normal growth. This is the case, for example, of the Antarctic bacterium *Pseudoalteromonas haloplanktis* TAC125, whose metabolic reconstruction embeds around 97% of the metabolic genes actually expressed during exponential phase (Fondi et al., 2014).

Since metabolic reconstructions are usually homology-based, classic metabolic reconstructions fail in identifying characteristic metabolic feature and/or pathways of microorganisms that are phylogenetically and/or functionally different from well-characterized ones. -omics integration strategies that we have illustrated in previous sections are a promising approach in this context, since the combination of heterogeneous information layers may contribute to identify and properly fill metabolic gaps left in reconstructions derived from sequence-based annotations. In this context, it is to be noted that software for exploiting –omics information during constraints-based modelling are mostly focused on the integration of transcriptomics (expression) data (Table 2). Accordingly, more effort is needed in the future to allow including other –omics datasets (e.g. proteomics, metabolomics etc.) when modelling microbial metabolism. Nevertheless, experiments with fine scale observation (e.g. single knock-out mutants) are usually decisive in providing a robust answer to knowledge gaps in homology-based reconstructed metabolic models.

Finally, taxonomic information is another, still not fully exploited, resource in metabolic reconstruction and very few works describe possible approaches for merging information from models of two (or more) close organisms. Nevertheless as the number of available and, most importantly, highly curated and experimentally validated metabolic models will increase and span over a larger phylogenetic range, this resource is expected to gain an important role. This is also relevant for experimental design and strains selection, providing insights into the issue of which species/genera should be sequenced with higher priority in order to increase the coverage of available metabolic information, Some works have tried quantifying the increase in information content when adding novel genomes to a given dataset and, importantly, the influence of phylogenetic distance among species in guiding this choice (“where to add taxa”) (Eddy, 2005; McAuliffe et al., 2005; Pardi and Goldman, 2005; Geuten et al., 2007). Similar approaches may thus be useful in the context of microbial metabolic biodiversity exploration, leading, for example, to the identification of novel, biotechnologically relevant pathways. Indeed, we underline that a thorough exploration of metabolic diversity may be obtained integrating taxonomic and metabolic information, guiding strains selection for more focused downstream postgenomics analyses.

## Conclusions

Many possible pathways link genotype and phenotypes, being represented by distinct functional states of cellular components (genes, metabolites, DNA methylation states, proteins). In an ideal future scenario, most of the information layers herein described and accounting for those functional states will be known for a given bacterium in different environmental conditions. In this situation, computational biology will be able to guide *in silico* reverse engineering and trace back the optimal path towards the desired microbial phenotype(s).

In this work we have reviewed some of these information layers and their growing exploitation in present-day microbiology research due to the spreading of massive –omics technologies. Each of them is bringing valuable insights into the comprehension of cellular architecture and functioning. Nevertheless, mounting evidences suggest that it is the integration of these large, multi-scale datasets that will bring alternative perspectives to our current view of microbial cell organization and possibilities for explain the emergence of complex phenotypes. Also, connections among biological features that were previously thought to be unrelated (e.g. epigenetics and pathogenicity) are being discovered following this path. The capability of integrating and combining multi –omics data with metabolic modelling techniques is expected to provide even more accurate and realistic predictions of cellular metabolism in those cases in which, for example, metabolic activity is decoupled from expression of the corresponding genes. In this context, genetic constraints (e.g. CAI) and epigenetics mechanisms (e.g. DNA methylation, post-translational modifications) represent still unexploited layers in metabolic modelling and a challenge for future *in silico* simulations.

## Acknowledgments

**During the realization of this work** Marco Fondi was supported by a FEMS advanced fellowship (FAF2012). This work is also supported by a PNRA (Piano Nazionale per la Ricerca in Antartide) grant (PNRA 2013/B4.02).

